# A catalytic-independent function of human DNA polymerase Kappa controls the pool of the Checkpoint Kinase 1

**DOI:** 10.1101/2021.02.09.430494

**Authors:** Marina Dall’Osto, Laura Pierini, Nicolas Valery, Jean-Sébastien Hoffmann, Marie-jeanne Pillaire

## Abstract

DNA polymerase kappa (Pol κ) has been well documented thus far for its specialized DNA synthesis activity during translesion replication, progression of replication forks through regions difficult to replicate and replication checkpoint at stalled forks.

Here we unveiled an unexpected role for Pol κ in controlling the stability and abundance of Chk1, the major mediator of the replication checkpoint. We found that loss of Pol κ decreased the Chk1 protein level in the nucleus of four human cell lines. Pol κ and not the other Y‐family polymerase members is required to maintain the Chk1 protein pool all along the cell cycle. We showed that Pol κ depletion affected the protein stability of Chk1 and protected it from proteasome degradation and the replication recovery defects observed in Pol κ-depleted cells could be overcome by the re-expression of Chk1. Importantly, this new function of Pol κ does not require its catalytic activity, revealing that in addition to its known roles in the replication process, Pol κ can contribute to the maintenance of genome stability independently of its DNA synthesis activity.

## INTRODUCTION

Cells have to continuously cope with a variety of DNA damages induced by the exposure to exogenous and endogenous genotoxic agents. Cellular responses such as signalling, repairing or bypassing the damage are required to deal with DNA damages and avoid the replication fork blockage during S-phase. Cells need to limit the accumulation of stalled forks, a process described as replication stress, in order to restrain transmission of DNA damages to daughter cells (Zeman & Cimprich, 2014; Técher *et al*, 2017). An important part of the cellular response to replication stress is the induction of the ATR/Chk1 replication checkpoint which senses stalled replication forks, allows their stabilization and repair, prevents the firing of late replication origins, and inhibits entry into mitosis until the completion of replication (González Besteiro & Gottifredi, 2015; Zhang & Hunter, 2014).

The ATR-Chk1 signaling axis is now described as a tunable brake required during unperturbed cell proliferation to couple the replication of the genome and the cell cycle progression (Panagopoulos & Altmeyer, 2020). ATR and Chk1 are essential actors in the genome stability maintenance. Indeed, mutations in *ATR* are responsible for an autosomal recessive disorder called Seckel syndrome (Lecona & Fernandez-Capetillo, 2018) and Chk1 heterozygosity leads to defects in cell cycle control, accumulation of DNA damages and predisposes cells to cancer (Lam *et al*, 2004). Under physiological conditions, the depletion of Chk1 decreases the global rate of replication (Petermann *et al*, 2006, 2010; Técher *et al*, 2016). To ensure correct connections between replication and cell cycle progression, the abundance of Chk1 that relies on its stability is critical for cellular stress response and checkpoint maintenance (Panagopoulos & Altmeyer, 2020). One of the most documented modes of regulation of Chk1 is the ubiquitination mediated by the Cullin-Ring E3-ubiquitin ligases Cul1, Cul4A, HUWE1 which ubiquitinate the C-terminal degron like region of Chk1 and target it for proteasomal degradation (Zhang *et al*, 2009, 2005; Cassidy *et al*, 2020). This ubiquitination-dependent regulation of Chk1 can be directly antagonized by the ubiquitin hydrolases USP1, USP3, USP7 and Ataxin3 which deubiquitinate Chk1 (Cheng & Shieh, 2018; Guervilly *et al*, 2011; Tu *et al*, 2017; Alonso-de Vega *et al*, 2014).

The translesion synthesis (TLS) is also an important mechanism to respond to replicative stress. It involves the translesional Y-family DNA polymerases Pol η, Pol κ, Pol i and Rev1 also called specialized polymerases. They facilitate the bypass of DNA lesions that block the replicative DNA polymerases by insertion of nucleotides opposite DNA lesions. They are devoid from exonuclease activity and show a flexible catalytic site able to adapt to damaged DNA template (Guo *et al*, 2009). Y-DNA polymerases are also implicated in DNA repair mechanisms such nucleotide excision repair or homologous recombination (McVey *et al*, 2016; Ogi *et al*, 2010; Lange *et al*, 2011). In addition to their TLS or repair functions, these specialized DNA polymerases also play critical roles in the replication of non B-DNA and in the prevention of replication stress induced by oncogenes (Bournique *et al*, 2018; Tsao & Eckert, 2018; Pillaire *et al*, 2014; Tonzi & Huang, 2019). For instance Pol η contributes to the stability of Common Fragile Sites (CFS) which have the potential to adopt non-B DNA structures (Bergoglio *et al*, 2013). The deoxycytidyl transferase Rev1 is required to replicate DNA sequences prone to form G4 secondary structures or enriched in nucleotide repeats (Barnes *et al*, 2017; Walsh *et al*, 2013). Similarly, Pol κ is required for the replication of structured DNA sequences and rescues the replicative DNA polymerase Pol δ when it is stalled at repetitive sequences within CFS (Sarkies *et al*, 2010). Pol κ is also needed to maintain viability upon replication stress induced by oncogene activation (Yang *et al*, 2017) since it is required to protect and to restart the replication forks following starvation of dNTPs (Tonzi *et al*, 2018).

Thus far, all these known Pol κ functional roles, i.e. TLS, replication of non-B structured DNA and repetitive sequences, DNA synthesis on unwound DNA at stalled forks were entirely associated to its DNA polymerase catalytic activity. Indeed, mutation of the residues D198 and D199 that belong to the catalytic site of Pol κ (Lone *et al*, 2007) abolished the capacity of the polymerase to extend primers *in vitro* (Ohashi *et al*, 2000), sensitized human cells to benzo[a]pyrene diolepoxide, mitomycin C and bleomycin (Kanemaru *et al*, 2015), decreased the repair of interstrand crosslinks (Williams *et al*, 2012) or impeded primer synthesis (Bétous *et al*, 2013) and fork restart (Tonzi *et al*, 2018) at stalled forks.

The multiple functions of Pol κ imply that its cellular level needs to be tightly regulated to avoid perturbation of genome maintenance. In untreated cells, its aberrant recruitment to replication forks in cells depleted for USP1 or p21CDNKN1 leads to a decrease of the fork speed and its overexpression is associated with an instability of CFS, DNA breaks and tumorigenesis in mice (Bavoux *et al*, 2005; Mansilla *et al*, 2016; Pillaire *et al*, 2007). In addition, dysregulation of Pol κ expression can affect its normal subcellular localization and can contribute to drug resistance (Peng *et al*, 2016; Temprine *et al*, 2020). Pol κ depletion in untreated cells induces hallmarks of replication stress with RPA foci formation and γ-H2AX in S phase which are indicative of endogenous ssDNA accumulation and DNA breaks respectively and common fragile site expression (Bétous *et al*, 2013; Mansilla *et al*, 2016) features also observed in absence of Chk1 (Gagou *et al*, 2010).

Here we unveiled an unexpected role for Pol κ in controlling the stability of Chk1 in mammalian cells. We found that depletion of Pol κ and not the other Y‐family polymerase members, induces a decrease of Chk1 protein level in the nucleus of four different human cell lines. Strikingly, this regulation is independent of the catalytic activity of Pol κ and occurs all along the cell cycle. Pol κ depletion does not affect the mRNA expression of Chk1 but favors its degradation through the proteasomal pathway. Finally, we found that the replication defects observed in Pol κ-depleted cells is linked to the low abundance of Chk1. Collectively, this work highlights a catalytic-independent function of Pol κ in genome maintenance.

## RESULTS

### 1. Chk1 protein level is reduced in mammalian cells depleted for Pol Kappa

To better understand the implication of the DNA polymerase Kappa (Pol κ) in the maintenance of genome stability, we depleted Pol κ from different human cell lines and analyzed the Chk1 signaling pathway. We found a decrease of Chk1 protein level in the nuclear fraction of MRC5-SV Pol κ depleted cells (Fig 1A to 1C) whereas the level of other proteins implicated in fork progression and ATR/Chk1 pathway such as claspin, TopBP1, Rad18, Rad17, Rad9A or Pol delta were not reduced (Suppl Fig 1A). As a control we treated cells with hydroxyurea, an inhibitor of ribonucleotide reductase which induces the activation of the ATR/Chk1 and the phosphorylation of Chk1 before its down-regulation that can be observed later on (Leung-Pineda *et al*, 2009). Analysis of Pol κ and Chk1 protein levels in different nuclear extracts of MRC5-SV showed a good correlation (R^2^=0.906, Fig 1B). To confirm this observation, we used immunofluorescence microscopy as a second approach. Chk1 fluorescence intensity was significantly decreased after Chk1 or Pol κ depletion in the nuclei of MRC5-SV cells (Fig 1C) and importantly the Chk1 protein level was rescued by ectopic expression of the polymerase (Fig 1C and suppl Fig 1C). Thus we revealed here that Pol κ depletion leads reproducibly to a decrease in Chk1 abundance in the nucleus, an observation that was not reported in previous studies using whole cell extracts from Pol κ depleted cells (Bétous *et al*, 2013; Mansilla *et al*, 2016; Tonzi *et al*, 2018). We verified that indeed there was no correlation between Chk1 and Pol κ protein levels in whole cell extracts (R^2^=0.002, Suppl Fig 1) as compared to nuclear extracts (R^2^=0.906, Fig 1B), supporting that the impact on Chk1 level occurs reproducibly in the nuclear compartment. We next checked whether this effect was also observed in additional cell lines. The fluorescence intensity of Chk1 was monitored in more than 500 nucleus of HCT116 and RKO colon cancer cells after depletion of Pol κ with or without reintroduction of GFP-Pol κ (Fig 1D, 1F and suppl 1D). Transfection with siRNA targeting Chk1 was used as a control of down expression of Chk1 (Fig 1D and 1F). The fluorescence intensity of Chk1 decreased by 37% and 28% respectively in HCT116 and RKO transfected with a siRNA targeting the coding sequence of *POLK* (Fig 1E and 1G) and by 27% in HCT116 transfected with siRNA targeting the 3’UTR of *POLK* (Fig 1E). Chk1 protein level was rescued by ectopic expression of Pol κ in HCT116 cells (Fig 1D, 1E and suppl Fig 1D). Western blots analysis also confirmed the results in HCT116, RKO and in a human embryonic kidney cell line (293T) (suppl Fig 1E). As in MRC5-SV cells, HU treatment induced a Chk1 protein decrease in the nucleus of 293T cells that is amplified by Pol κ depletion (0.5 versus 0.1 relative quantity of Chk1 in control and Pol κ depleted cells respectively) (suppl Fig 1E). Collectively, these data support the notion that Pol κ is required to maintain the Chk1 protein level in the nucleus of human cells.

**Figure 1:**
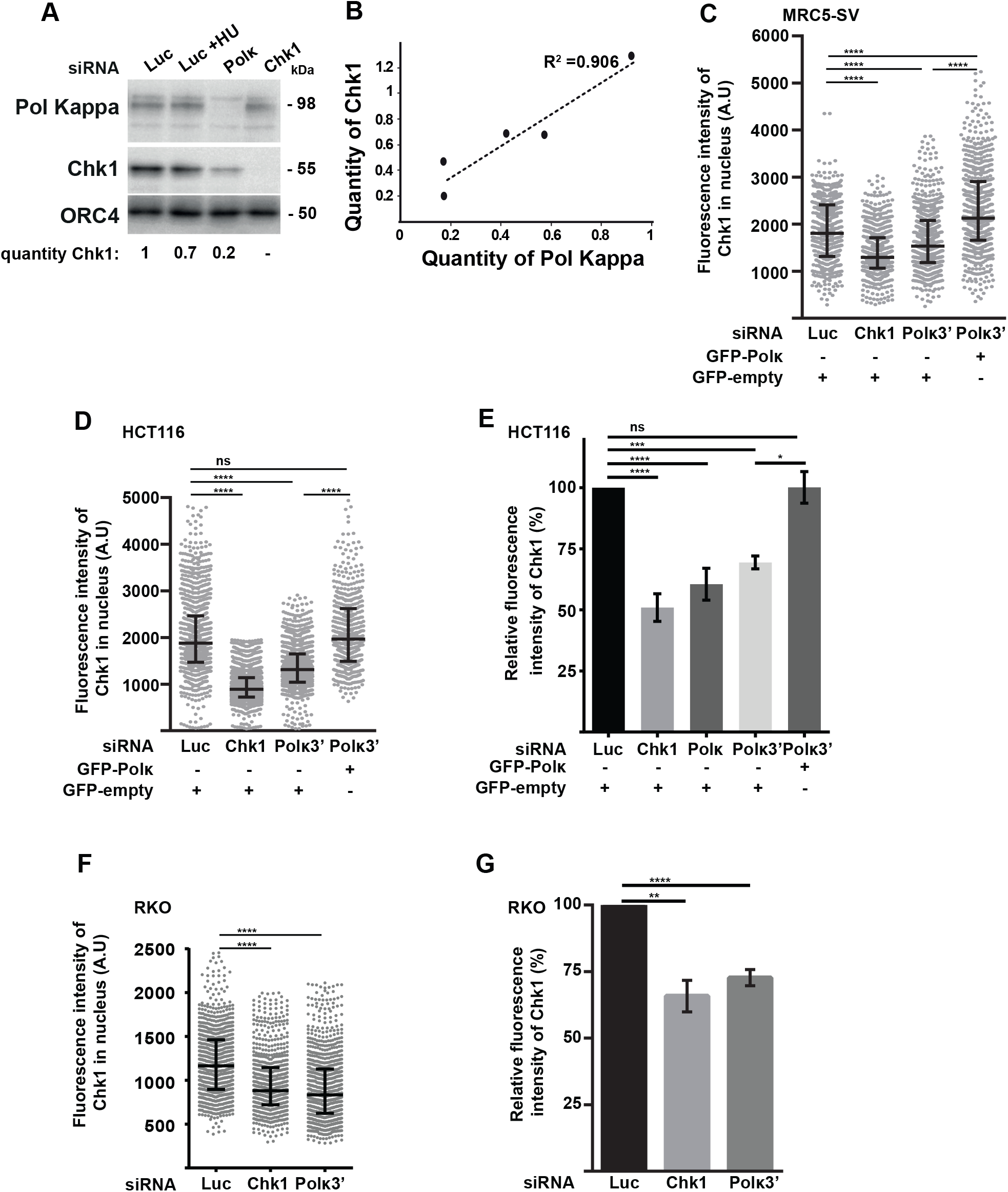
Chk1 protein level is reduced in the nucleus of mammalian cells depleted for Pol κ. (A) Western blot analysis of Chk1 and Pol κ in MRC5-SV nuclear extracts 48h after transfection with a control (Luc), Pol κ (Polκ) or Chk1 (Chk1) siRNA. Cells were untreated or treated with 1mM HU for 1h. ORC4 is shown as a protein-loading control. Quantification of Chk1 is relative to siLuc condition. (B) Relative Chk1 or Pol Kappa protein levels in siPolk nuclear extracts were normalized to siLuc condition (quantity of Chk1, quantity Pol of Kappa respectively) and data from 5 independent experiments in MRC-SV are plotted. The regression curve (dashed line) and R-square are shown. MRC5-SV (C), HCT116 (D) and RKO (F) cells were co-transfected with the indicated siRNAs and a vector expressing either GFP-empty or GFP-Pol κ. The fluorescence intensity of Chk1 was quantified in each nucleus. Medians with 25% and 75% interquartile ranges were represented. (***p=0.001; ****p<0.0001; Mann-Whitney test). A.U = arbitrary units. ns = not significant. Relative fluorescence intensity of Chk1 in nuclei of HCT116 (E) and RKO (G) cells transfected with the indicated siRNAs (G) or both with a vector expressing either GFP-Pol κ or GFP-empty (E). Values are the mean (± SEM) of medians of three or four independent experiments. Relative fluorescence intensity was adjusted to siLuc condition. (*p<0.05;**p<0.01; ****p<0.0001; t-test). ns = not significant.

### 2. Among the Y-DNA polymerase family, only the Polk depletion causes a Chk1 nuclear drop

To check whether the effect on Chk1 is specific to Pol κ, HCT116, RKO and 293T cells were transiently transfected with siRNA targeting the three other Y-family TLS DNA polymerases, Pol eta, iota and REV1, and the Chk1 fluorescence intensity was measured (Fig 2A and 2B, suppl Fig 2C). siRNA transfection efficiencies were concomitantly checked by RT-qPCR (suppl Fig 2A and 2B). While depletion of Chk1 itself, Pol κ or USP7, an ubiquitin hydrolase already shown to stabilize Chk1 (Alonso-de Vega et al., 2014) triggered a Chk1 drop in the nucleus, the depletion of Pol iota, eta or Rev1 did not, supporting that Chk1 is specifically regulated by Pol κ.

**Figure 2:**
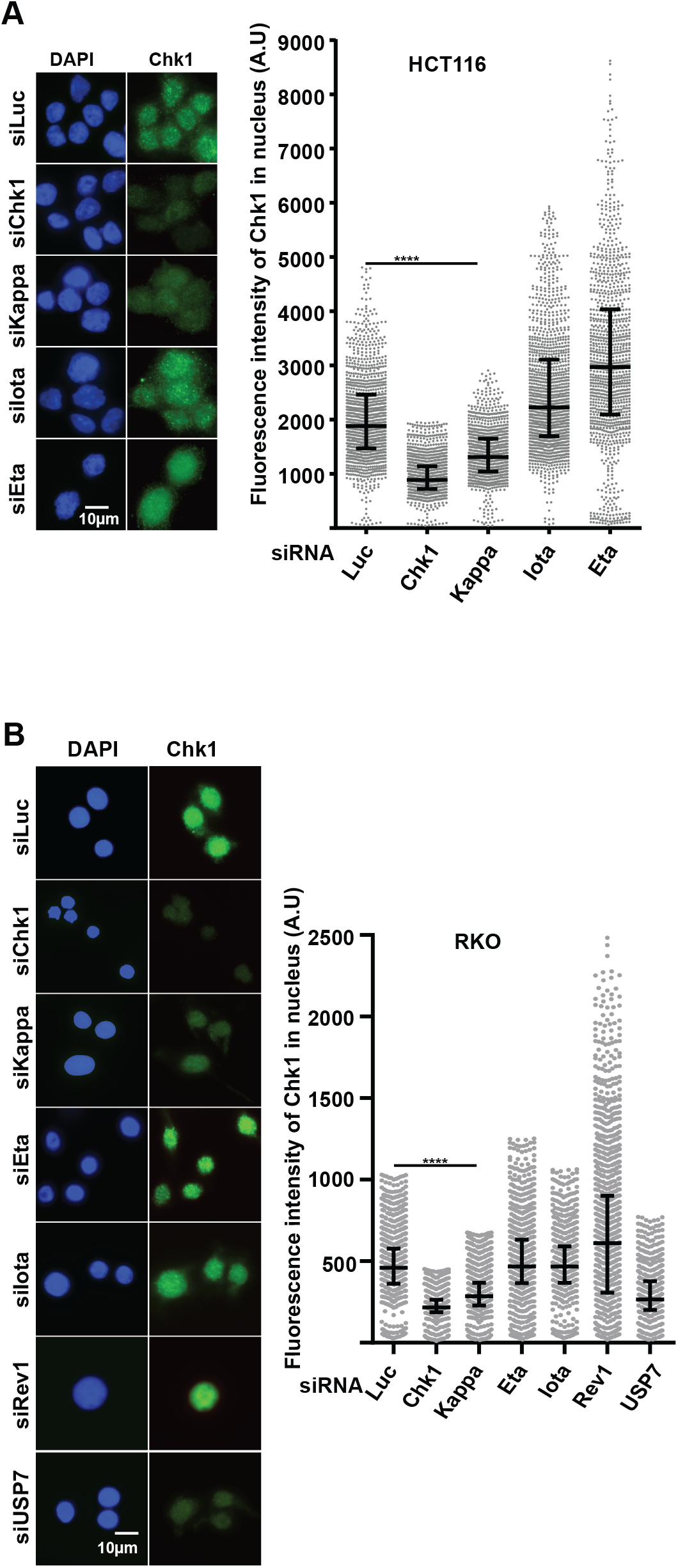
Among the Y-DNA polymerase family, only the Pol κ depletion causes a Chk1 nuclear drop. Representative images and quantification of Chk1 immunostaining (green). HCT116 (A) and RKO (B) cells were transfected with the indicated siRNAs and DNA was stained with DAPI. The fluorescence intensity of Chk1 was quantified in each nucleus. Medians with 25% and 75% interquartile ranges were presented. (****p<0.0001; Mann-Whitney test). Scale bar, 10 μm. A.U = arbitrary units.

### 3. The Pol κ dependent Chk1 downregulation is not due to cell cycle arrest in G1

Chk1 protein level in human cells has been shown to accumulate in S and G2 phases and display its lowest level in G1 (Kaneko *et al*, 1999). We confirmed these data as shown in suppl Fig 3A. To exclude the possibility that the Chk1 decrease in Pol κ depleted cells is the consequence of an enrichment in G1 population, we compared the cell cycle distribution of Pol κ depleted cells (siPolκ3’and siPolκ) with control mocked-depleted cells (siLuc) by Quantitative Image-Based Cytometry (QIBC, (Toledo *et al*, 2013)), an experimental approach which monitors the Chk1 fluorescence intensity all along the cell cycle in HCT116 cells and by FACS in RKO cells (suppl Fig 3B). We found that 30 to 38% of the Pol κ depleted cells versus 39% in controls were in G1 in HCT116 cells (suppl Fig 3B left panel) and 40% to 45% of the Pol κ depleted cells versus 43% in controls were in G1 in RKO (suppl Fig 3B right panel), demonstrating that no enrichment in G1 cell population occurred when the pool of Chk1 is affected in Pol κ-depleted HCT116 and RKO cells with no modification of the global cell cycle progression, in contrast to cells which display a total depletion of Chk1 expression (Fig 3B right panel). When we combined the quantification of Chk1 fluorescent intensity, EdU incorporation and DNA content (DAPI labeling) (Fig 3), we found that Pol κ depletion impacted Chk1 protein level in all the phases of the cell cycle. These observations suggest that Pol κ-dependent regulation of Chk1 does not result from a cell cycle modification.

**Figure 3:**
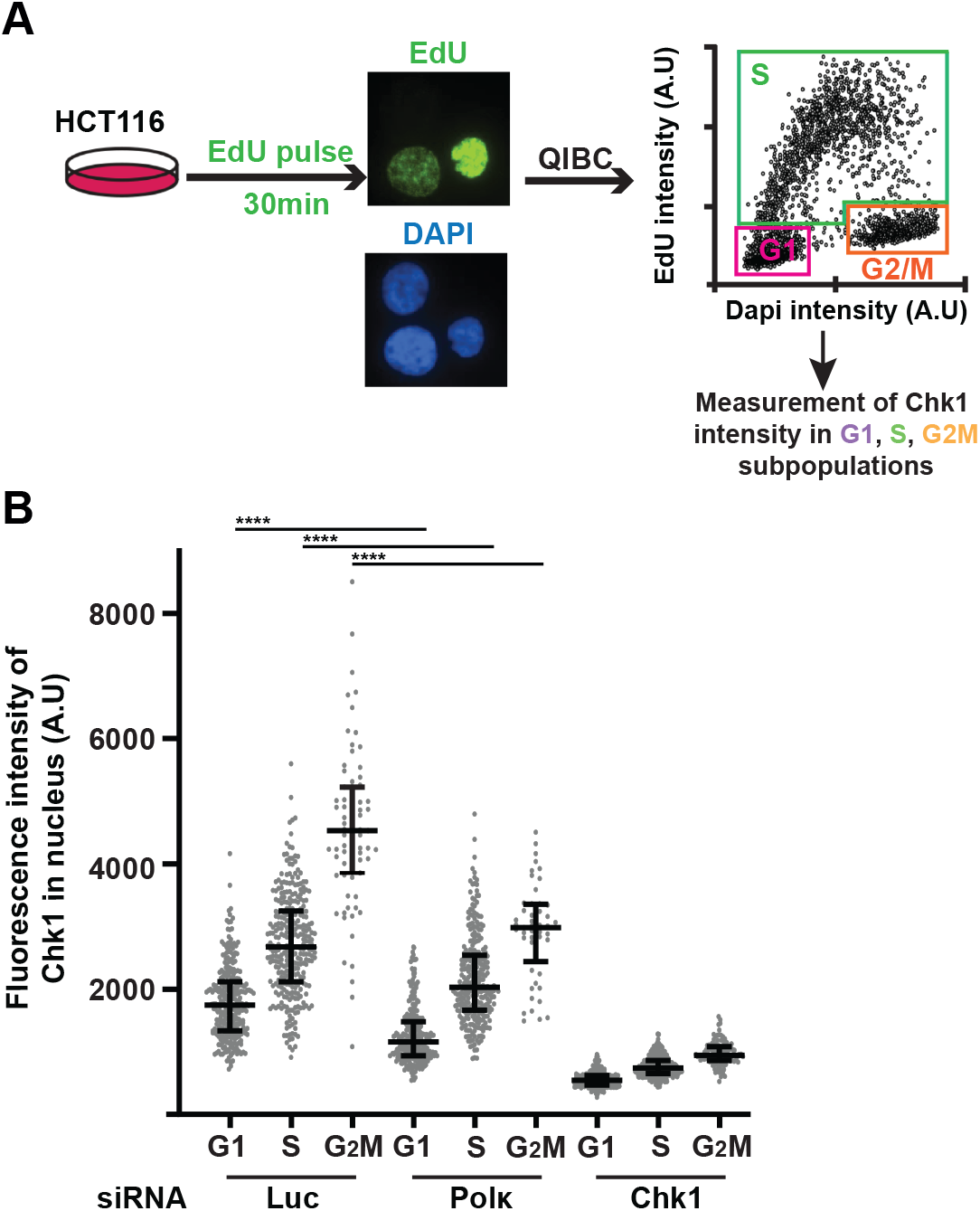
The Pol κ-dependent Chk1 downregulation is not due to cycle arrest in G1. (A) Schematic representation of pulse-EdU labelling experiment. HCT116 Cells were transfected with control (Luc), Pol κ (Polκ) or Chk1 (Chk1) siRNA. 48h after transfection, asynchronous cells were pulse-labelled with EdU for 30 min (10μM). Incorporated EdU was coupled to Alexa 488 in click-it reaction and DNA was stained with DAPI. EdU/DAPI dot plot shows the cell cycle distribution. (B) The fluorescence intensity of Chk1 was quantified in each nucleus of each cellular subpopulation transfected with indicated siRNA. Medians with 25% and 75% interquartile ranges were represented. A.U = arbitrary units.

### 4. Pol κ controls Chk1 abundance independently of its DNA synthesis activity

For the Pol κ functions such as replication of non-B structured DNA and repetitive sequences, bypass of DNA damages, full activation of the S-phase checkpoint, the catalytic activity of the polymerase is required. To determine whether this is also true for maintaining the level of Chk1, cells depleted for Pol κ with a 3’UTR *POLK* siRNA were transfected with a vector coding for the catalytically inactive form of Pol κ (GFP-Polk-Dead), and Chk1 level was quantified by immunofluorescence. The results show that, similarly to the WT Pol κ (Fig 1C to E), the Chk1 fluorescence intensity was also restored by expression of the Dead Pol κ in RKO (Fig 4A) and HCT116 (Fig 4B) cells, demonstrating that the polymerase activity of Pol κ is not required to maintain the Chk1 protein level in human cells.

**Figure 4:**
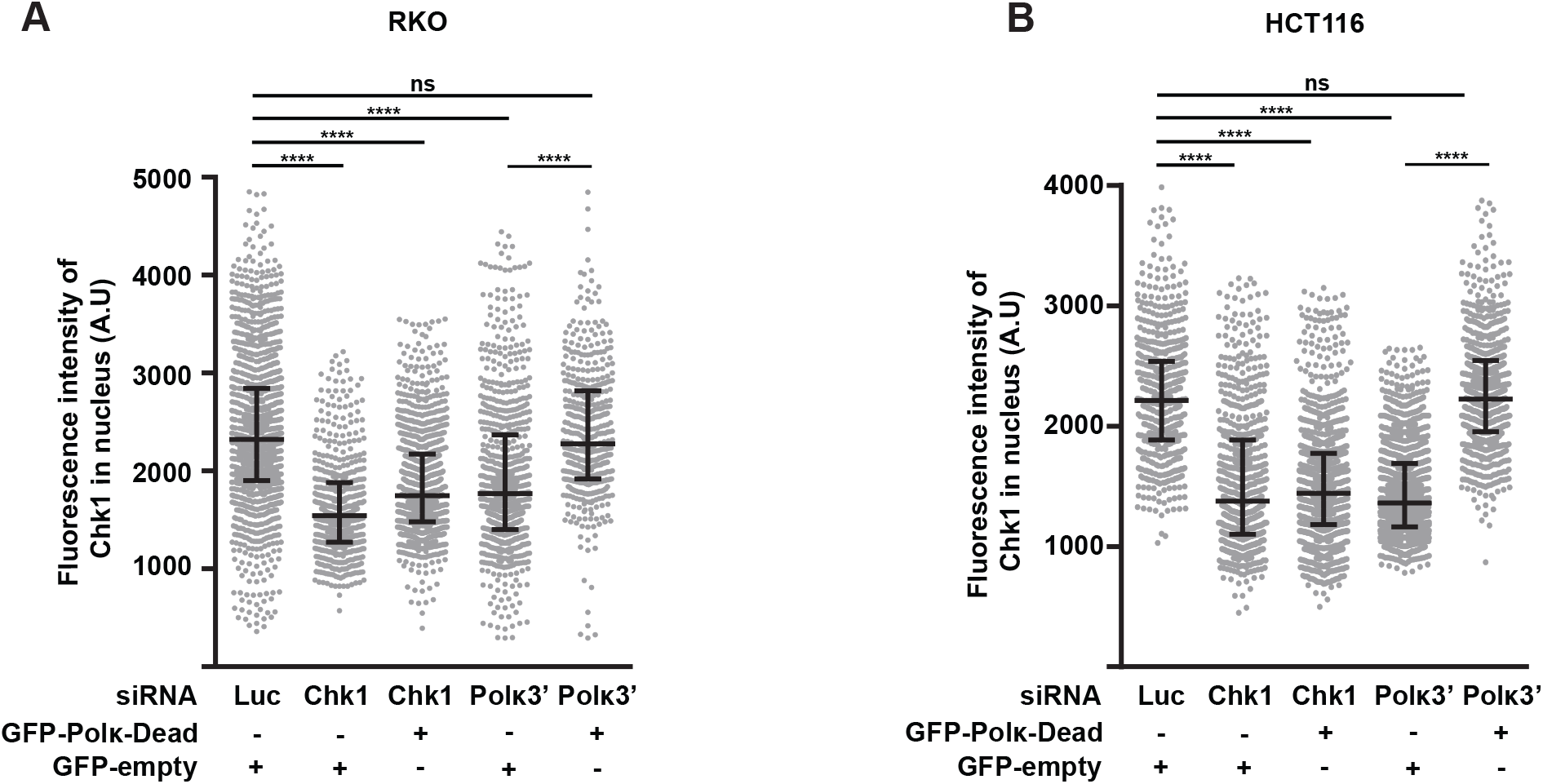
Pol κ controls the Chk1 pool independently of its DNA synthesis activity. Quantification of Chk1 immunostaining in RKO (A) and HCT116 (B) cells co-transfected with the indicated siRNAs and with a control vector expressing GFP-empty or GFP-Polκ-Dead. 48h after transfection, the fluorescence intensity of Chk1 was quantified in each nucleus. Medians with 25% and 75% interquartile ranges were represented (****p<0.0001; Mann-Whitney test). A.U = arbitrary units. ns = not significant.

### 5. Pol κ protects Chk1 from degradation

Next, we wanted to determine how Pol κ affects Chk1 expression. To deal with this question we carried out reverse transcription and real-time quantitative PCR to analyze the mRNA level of Chk1 in Pol κ depleted cells with two different siRNA. We did not find any significant difference when compared to control cells in four different cell lines (Fig 5A). This observation rules out any potential implication of Pol κ in *CHEK1* gene promoter repression, or its binding to the *CHEK1* transcript, to explain the down regulation of Chk1. These data are in accordance with the observations presented in Fig 4A and 4B showing that Pol κ re-expression could not restore Chk1 fluorescence induced by a Chk1 siRNA-mediated depletion. It has been reported that Chk1 can be targeted by ubiquitin ligases to control its stability and its degradation through the proteasome (Zhang *et al*, 2005, 2009; Cassidy *et al*, 2020). To test if Chk1 instability resulted from proteasomal degradation, we treated Pol κ depleted cells with the proteasome inhibitor MG132 to block the proteolysis and we monitored the level of Chk1 by immunoblotting (Fig 5B). The results clearly showed in three different cell lines that the proteasome inhibition lead to a stabilization of Chk1 in Pol κ depleted conditions. Treatment of HCT116 with cycloheximide (CHX), an inhibitor of protein synthesis, induced the decrease of Chk1 and p53 protein levels in the control condition and only Chk1 was further affected under Pol κ depletion (Fig 5C). Addition of MG132 restored Chk1 and p53 levels in CHX treated conditions. Collectively, these data support that the defective Chk1 pool in Pol κ depleted cells is not the consequence of a modification of its mRNA abundance but rather the result of its enhanced degradation by the proteasome, and thus Pol κ seems to be a requisite factor to stabilize Chk1 protein level in human cells.

**Figure 5:**
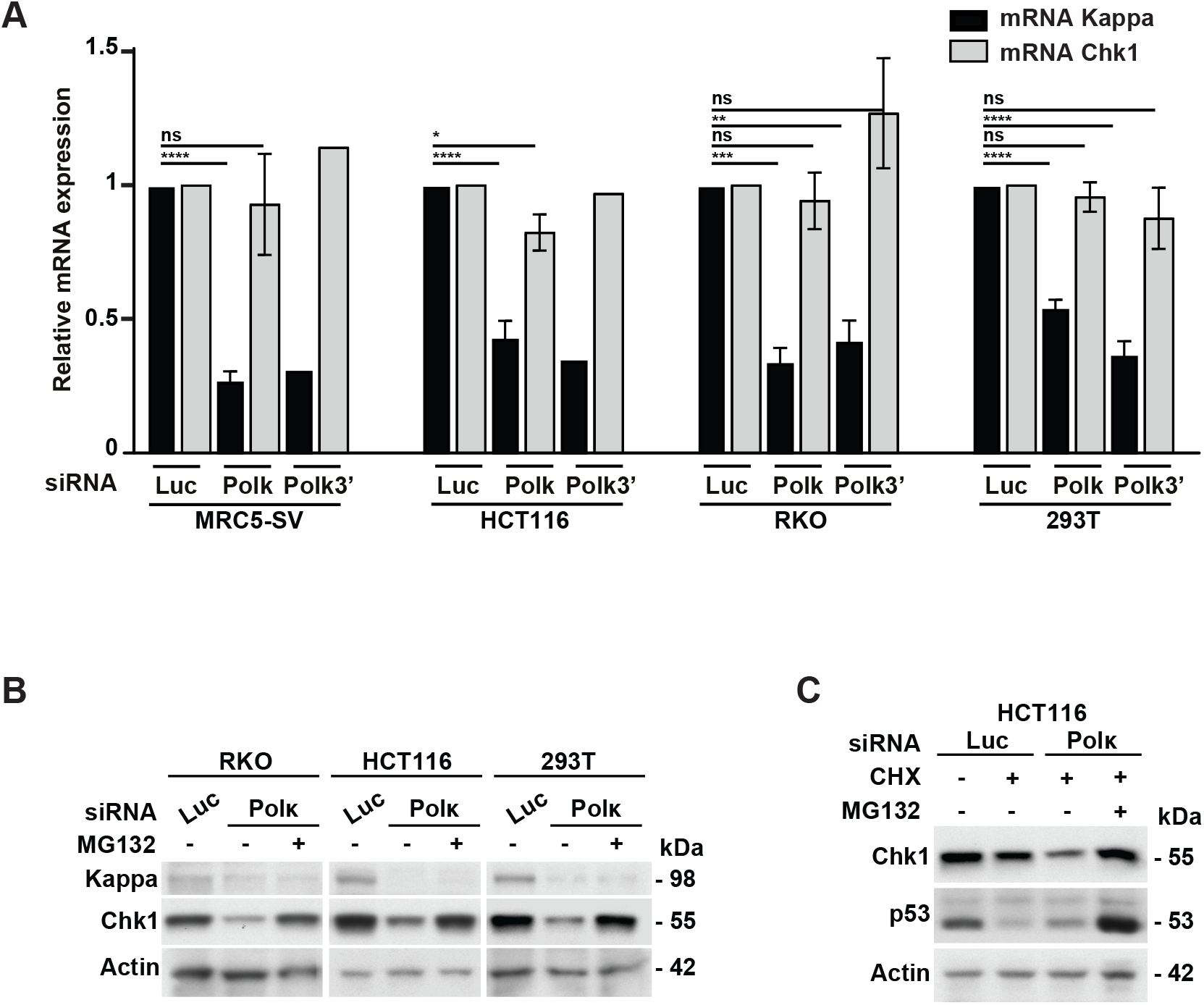
Pol κ protects Chk1 from degradation. (A) Transcripts levels analysis of Pol κ and Chk1 genes by RT-qPCR in MRC5-SV, HCT116, RKO and 293T cells transfected with siRNA against the 3’UTR of Pol κ (Polĸ3’), the coding sequence of Pol κ (Polκ). Relative expressions were normalized to siLuc condition. Values are the mean (± SEM) of medians of independent experiments. (*p<0.05; (**p<0.01; ***p<0.001;****p<0.0001; t-test) ns = not significant. (B) Western Blot analysis of Chk1 in RKO, HCT116 and 293T cells. 48h after transfection with a control siRNA (Luc) or Pol κ siRNA (Polκ), cells were treated or not with MG132 (20μM) for 4h just after transfection then 6h before to harvest. Actin is shown as a protein-loading control. (C) Western Blot analysis of Chk1 in HCT116 cells extracts. 48h after transfection with indicated siRNA cells were treated with 50 μg/mL of cycloheximide (CHX) in addition or not of MG132 (20μM) for 5h. Actin is shown as a protein-loading control.

### 6. The replication defects associated with Pol κ loss can be restored by Chk1 ectopic expression

Considering the fact that Chk1 is required for the global genomic replication in absence of exogenous stress (Petermann *et al*, 2006; González Besteiro *et al*, 2019), we asked whether the depletion of Pol κ can affect the genome replication to the same extent. We pulse-labeled cells with EdU and we measured the EdU intensity in the S-phase cell population as in figure 3. We found that the depletion of Pol κ decreased the EdU incorporation to a similar extent as the Chk1 depletion (Suppl Fig 4), meaning that Pol κ is required to maintain the global replication in absence of exogenous stress. Pol κ was shown to be required for restart of stalled fork after HU treatment (Tonzi *et al*, 2018). As Chk1 is also implicated in this process (Hromas *et al*, 2012; Smits & Gillespie, 2015), we wondered whether the fork restart deficiency observed by Tonzi *et al* in RPE Pol κ depleted cells can be explained by the attenuated abundance of Chk1 induced by the polymerase depletion. To test this hypothesis, we performed DNA fiber spreading analysis. Cells were labelled with IdU, treated with HU during 1h at 1mM, and released in fresh media containing CldU to label the restarting forks (Fig 6). The length of the CldU tracts is indicative of the replication recovery efficiency after fork stalling. As shown in Fig 6, we found that Pol κ depletion significantly reduced the track length by 44% (2.39 μm in siPolκ versus 4.23 μm in control cells) and reintroduction of GFP-Polκ in siPolκ cells restored the CldU track length. Similarly to Pol κ depleted cells, we observed that in Chk1 depleted cells the track length is reduced by 46% (2.27 μm in siChk1 versus 4.23 μm in control cells). Strikingly, we observed that the inefficient fork recovery in Pol κ-depleted cells was restored by expression of ectopic myc-Chk1, whereas the expression of GFP-Pol κ could not restore the fork recovery in Chk1-depleted cells. Importantly these data demonstrate that in addition to the role of Pol κ in the fork restart showed by Tonzi et al, the deficiency of the replication stress recovery observed in Pol κ depleted cells may rely on an insufficient pool of Chk1 in the nucleus.

**Figure 6:**
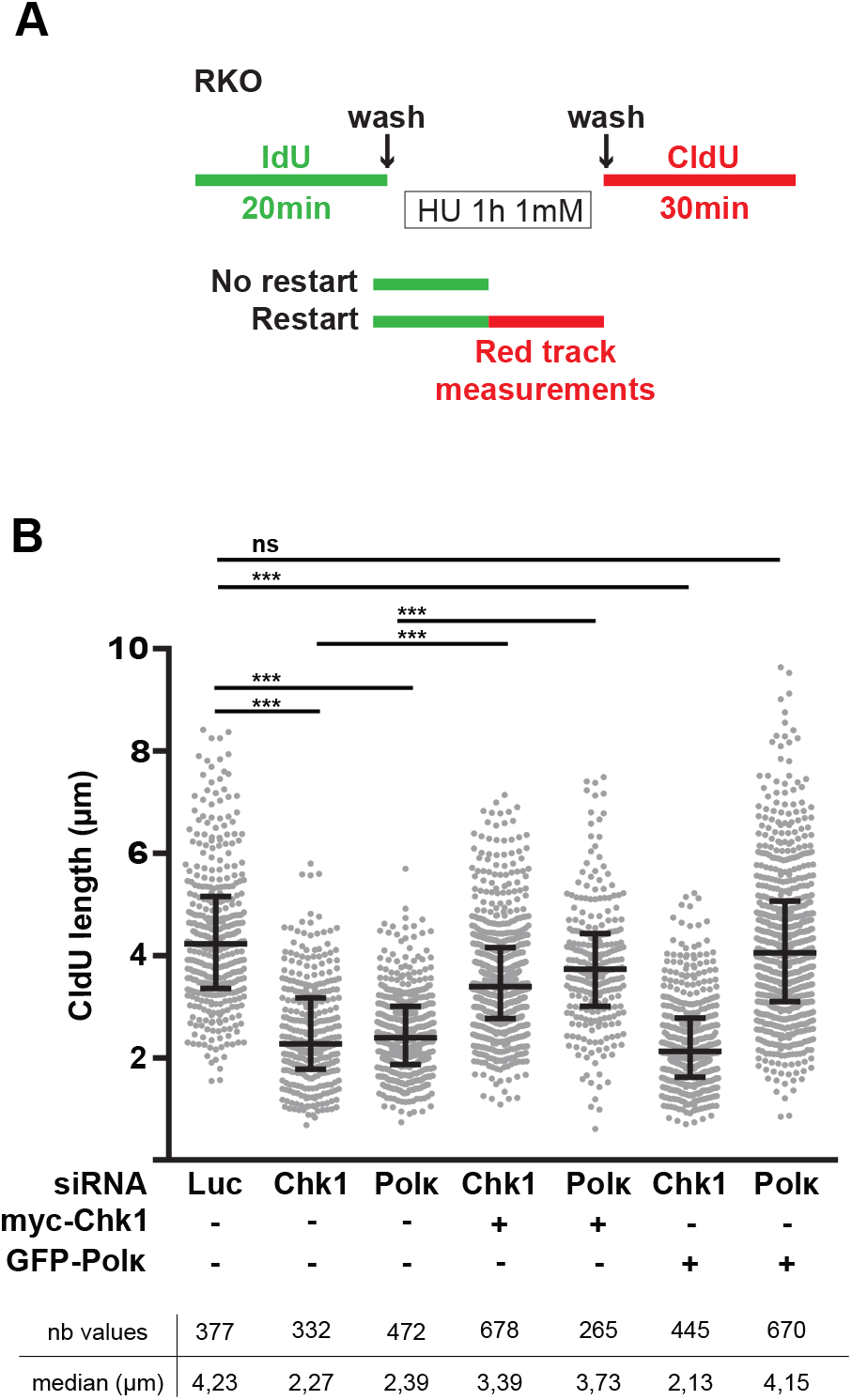
The replication defects associated with Pol κ loss can be restored by Chk1 ectopic expression. (A) Schematic representation of experimental DNA fiber labeling. RKO cells were co-transfected with the indicated siRNAs and a vector expressing either GFP-empty (-), GFP-Pol κ (GFP-Polκ) or myc-Chk1. 48h later, ongoing DNA replication forks were labeled with IdU (green tracks) for 20 min, treated with HU 1mM during 1h and then labelled with CldU (red tracks) during 30 min. (B) The length of CldU tracks were measured and the values of two independent experiments were pooled and plotted. The number of CldU tracks measured and the medians of their lengths are indicated in the table. Medians with 25*% and* 75*%* interquartile ranges are represented (***p=0.001; ***p<0.001; Mann-Whitney test). ns = not significant.

Collectively, all these findings highlight a non-catalytic function of Pol κ besides its well-documented TLS function to maintain proper DNA replication and ensure recovery from replication stress through the stability of Chk1.

## Discussion

In the absence of exogenous stress, the depletion of Pol κ is associated with increased rates of mutagenesis in mice, DNA breaks, sister-chromatid exchange, expression of common fragile site in human cells and increased number of 53BP1 nuclear bodies in G1 phase cells (Bétous *et al*, 2013; Hakura *et al*, 2019; Mansilla *et al*, 2016; Peng *et al*, 2016; Tonzi *et al*, 2018; Stancel *et al*, 2009). These hallmarks of genetic instabilities can reflect a direct role of Pol κ in the genomic replication. Indeed Pol κ is required to copy repetitive sequences known to be fork-stalling sites for replicative DNA polymerase delta (Barnes *et al*, 2017; Walsh *et al*, 2013), tolerate stress induced by oncogenes (Yang *et al*, 2017) and promote efficient fork progression under low level of dNTP (Tonzi *et al*, 2018). We have also previously shown that upon activation of the replication checkpoint with HU, Chk1 phosphorylation is dependent on the presence of Pol κ (Bétous *et al*, 2013) suggesting that Pol κ could be involved in the activation of the ATR-Chk1 axis. This has been further confirmed in human glioblastoma cell lines treated by the alkylating drug temozolomide (Peng *et al*, 2016). All the functions reported thus far have been demonstrated to require the catalytic DNA synthesis activity of the polymerase.

Here we present evidences supporting a new catalytic-independent role for Pol κ in the cellular homeostasis through the regulation of Chk1 abundance in human cells. We provide evidences that the depletion of Pol κ in four independent cell lines induces a decrease of the Chk1 protein level. It is specific to Pol κ since this Chk1 reduction is 1) rescued by ectopic GFP-Pol κ and 2) not shared by other DNA polymerases, consistent with previous reports where no modification of Chk1 protein level was observed in Pol η depleted or mutated cells (Despras *et al*, 2010). *CHK1* knock-out is lethal in mice and Chk1 haploinsufficiency leads to carcinogenesis (Liu *et al*, 2000). In contrast, the effects of Chk1 instability in Pol κ knock-out mice are milder with increased mutagenesis without affecting viability and with no cancer incidence (Stancel et al., 2009), suggesting that low level of Chk1 is enough to maintain genome stability and viability. Interestingly, Gonzalez-Besteiro and colleagues reported recently that indeed in Chk1-depleted cells, low levels of exogenous Chk1 was sufficient to rescue origin firing and restore DNA damage signalling (González Besteiro *et al*, 2019).

The level of Chk1 in the nucleus depends on a tight equilibrium between synthesis, degradation and nuclear export, and an excess or a lack of Chk1 can be deleterious for the genome stability (Smits & Gillespie, 2015). Indeed a higher abundance of Chk1 protein level restricts the replication stress induced by oncogenes or therapeutic treatments and contributes to malignant transformation (David *et al*, 2016; López-Contreras *et al*, 2012). Conversely, *CHK1* deficiency is associated with modifications of replication dynamics, mitotic defects, transmission of under-replicated DNA, and predisposition to cancer (Bétous *et al*, 2013; Petermann *et al*, 2006, 2010; Speroni *et al*, 2012; Técher *et al*, 2016; Lam *et al*, 2004). The tumor suppressor p53 has been shown to regulate the mRNA level of Chk1 and to downregulate its expression in response to stress signals (Gottifredi *et al*, 2001). Our data support that Pol κ effect on Chk1 is independent of p53 since 1) the mRNA of Chk1 is unchanged in Pol κ depleted cells, 2) the reduction of Chk1 is observed in p53 proficient cells (RKO, HCT116) as well as p53 deficient cells (293T and MRC5-SV). We also showed that the Chk1 downregulation cannot be the consequence of a cell cycle effect induced by the depletion of Pol κ and Pol κ down-regulate Chk1 all along the cell cycle.

Based on our results, we propose that the Chk1 decrease in Pol κ depleted cells can be the consequence of a lack of protection against its normal degradation. We observed a faster rate of Chk1 disappearance in Pol κ depleted cells that can be antagonized by proteasomal inhibition. We propose that Pol κ is a regulator of Chk1 stability as it was already observed for Rad17, another member of the ATR-Chk1 signaling pathway. However how Pol κ protects Chk1 from degradation remains to be determined.

Interestingly we showed that the catalytic activity of Pol κ is not required to maintain the Chk1 protein level, as the GFP-Pol κ dead also restored the Chk1 protein level in cells depleted for endogenous Pol κ. Non-catalytic function of other polymerases has been previously proposed. Pol η was shown to act as a bridge between PCNA and the ubiquitin-protein ligase Rad18, and to favor the mono-ubiquitination of PCNA by Rad18 independently of its polymerase activity (Durando *et al*, 2013). The c-ter of Rev1 but not the catalytic activity, is necessary to recruit the TLS DNA polymerases η κ i and ζ for DNA damage tolerance (Tissier *et al*, 2004). The Chk1-regulatory function of Pol κ which is independent of its catalytic activity could help in understanding some recent observations from the literature: Human lymphoblastic Nalm6 cells engineered to express a catalytic dead Pol κ mutant showed the same sensitivity to oxidative stress induced by hydrogen peroxide and menadione as Pol κ wild type cells while Pol κ knock-out counterpart were highly sensitive (Kanemaru *et al*, 2015). Temprine and colleagues have associated the increased expression of Pol κ to drug resistance but without a high rate of mutagenesis. They propose a non-catalytic function of Pol κ to explain the drug resistance (Temprine *et al*, 2020). Pol κ can also protect forks against nascent DNA degradation and the polymerase activity is not essential to perform this task (Tonzi *et al*, 2018). We propose that the stabilization by Pol κ of Chk1 could explain both the recovery of the stalled fork, the prevention of fork degradation and breaks accumulation (Forment *et al*, 2011; Murfuni *et al*, 2013; Thompson *et al*, 2012).

The maintenance of a basal replication checkpoint activity in unchallenged cells is an important concept that is reinforced with recent publications (Panagopoulos & Altmeyer, 2020; Michelena *et al*, 2019). The idea presented by Panagopoulos and Altmeyer is that facing endogenous replication stress due to local impediments such as repeats, structured DNA or local dNTP pool decrease, cells adapt instead of arrest their cell cycle progression. They proposed a fine-tuned deceleration and brake release mechanism dependent on the ATR-Chk1 axis (Panagopoulos & Altmeyer, 2020). Thus a constant basal activity of Chk1 is pre-requisite and Pol κ could be an actor of this regulation as in response to huge replication stress the dependency on Pol κ to restart stalled forks is less obvious (Tonzi *et al*, 2018).

Thus Pol κ could act at stalled forks to maintain genome stability in several ways: insertion of nucleotides in front of lesions (TLS function), elongation of primers to allow the S-phase checkpoint activation (S-phase checkpoint associated function), and regulation of the abundance of Chk1 in the nucleus (Chk1 regulator). The first two roles depend on its polymerase activity whereas the third one does not (Fig 7).

**Figure 7:**
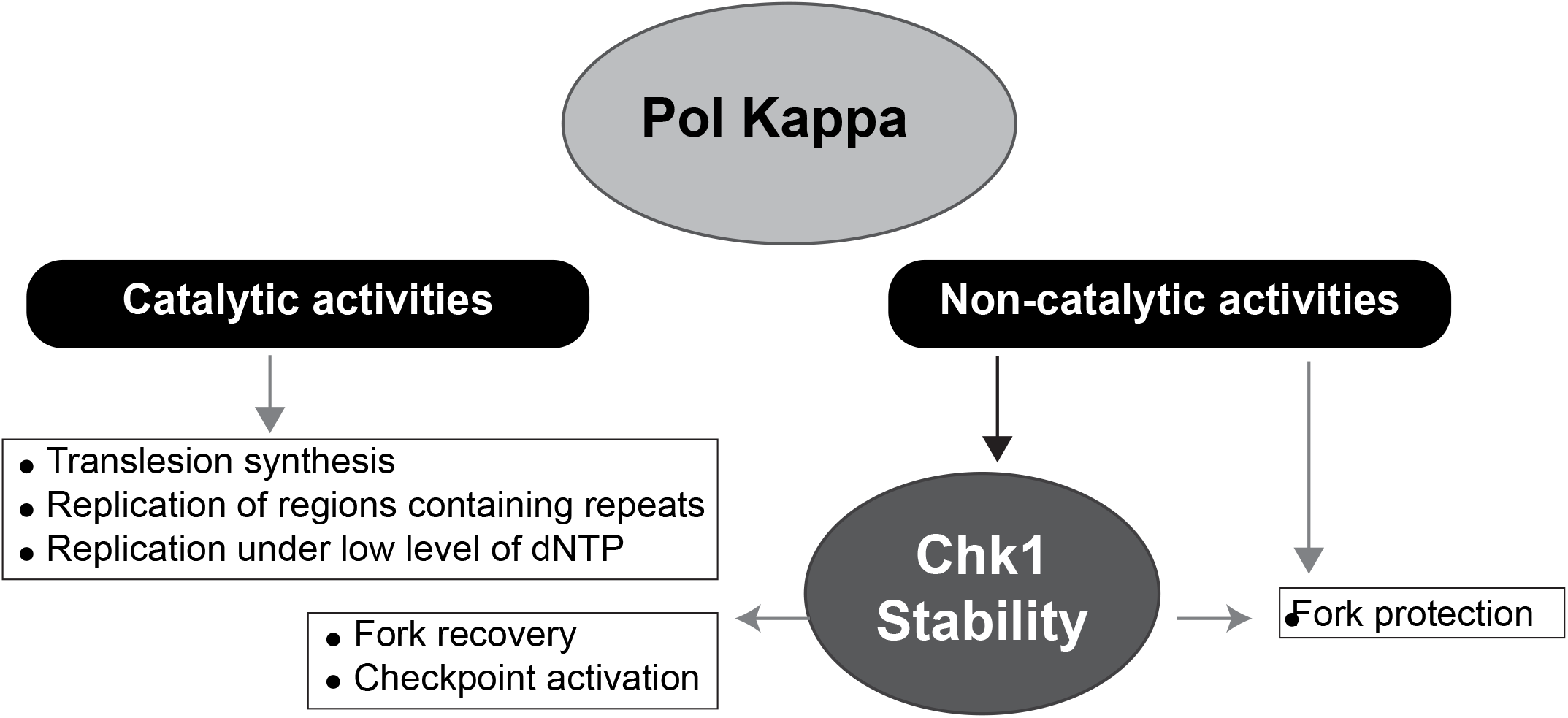
Working model. Under physiological conditions, the DNA polymerase activity of Pol κ is required to perform translesion synthesis, to replicate regions of the genome that contain repetitive sequences or local dNTP imbalance, to overcome fork stalling due to endogenous impediments and to participate in the S-phase checkpoint activation. In addition, the non-catalytic function of Pol κ may be fundamental to protect forks from degradation and to stabilize Chk1 which in turn acts to coordinate the response to fork stalling.

In conclusion, besides the well-documented importance of the catalytic function of the DNA Pol κ for translesion synthesis, stalled forks recovery, checkpoint activation or replication under low level of dNTP, this study unveils a unprecedented described DNA synthesis-independent regulatory function of Pol κ to protect forks from degradation and maintain basal level activity of replication checkpoint.

## Materials and methods

### Cell culture and cell lines

RKO, 293T, HCT116 Cells were cultured in DMEM/Glutamax supplemented 10% FBS (GIBCO) and MRC5-SV in alpha-MEM /Glutamax with 10% SVF (GIBCO) at 37°C in a humidified incubator in an atmosphere containing 5% CO_2_ (Panasonic MCO-19AIC-PE). All cells were routinely checked for mycoplasma contamination using PlasmoTest™ Kit (InvivoGen).

### Drugs and cell culture supplement

When indicated, cells were treated with hydroxyurea (Sigma-Aldrich), proteasome inhibitor MG132 (APExBIO), cycloheximide (Biosciences) for doses and time indicated in the figure legends. The dNTP analogs CldU and IdU (Sigma-Aldrich) were used as indicated in figures and the “DNA fibers assay” section of Materials and Methods.

### siRNA and plasmid transfections

1.5-2 × 10^6^ cells were seeded 24h before transfection with 45 nM of siRNA using Lipofectamine RNAiMAX (Life Technologies). The following siRNA molecules used against the coding sequence of Pol κ (Polκ) (5’CCAAUAGACAAGCUGUGAU3’ from Sigma-Aldrich), the 3’UTR of Pol κ (Polκ3’) (5’ACUCCAGCCUGAAGAGCGA3’ from Sigma-Aldrich), USP7 (5’CCCAAAUUAUUCCGCGGCAAA3’ from Sigma-Aldrich), the coding sequence of Chk1 (5’GAAGCAGUCGCAGUGAAGA3’ from Sigma-Aldrich), the 3’UTR of Chk1 (3’Chk1) (5’CUGGUGAAUAUAGUGCUGCUA3’ from Sigma-Aldrich), Polι (SMART pool: 5’CCACAGUUGGUAUUAGUUA3’; 5’GCACUAUGGUCGUGAGAGU3’; 5’CGGGUCAUGUAUACAAUAA3’; 5’GAACAUCAGGCUUUAAUAG3’ from Dharmacon), the 3’UTR of Polη (5’GCAAUGAGGGCCUUGAACA3’ from sigma-Aldrich) and Rev1 (5’CAGCGCAUCUGUGCCAAAGAA3’ from Sigma-Aldrich). For control, siRNA against luciferase and the catalytically inactive mutant of human Pol κ (Dead Pol κ - D198A and E199A) were previously described in Betous *et al*. (Bétous *et al*, 2013). The wild-type human Pol κ coding sequence was cloned into the peGFP vector (peGFP-Kappa_WT). Transfections of 4 to 8 μg of plasmids were carried out using JetPrime (Polyplus transfection) following the manufacturer’s recommendations.

### Nuclear cell extracts, Western Blot and quantification

Nuclear extracts were obtained using to the NE-PER™ Nuclear and cytoplasmic Extraction reagents kit (Thermo Scientific) according to the manufacturer’s recommendations. Proteins were dosed by the Bradford’s method (BioRad) and separated by SDS-PAGE (life technologies, NUPAGE™ 4-12% or 3-8%). For immunoblotting, primary antibodies were incubated overnight at 4°C in TBS 1X with Tween 0.1%. Secondary Peroxidase-coupled antibodies (Jackson Immuno Research, Life Technologies) were incubated at room temperature for 1 hr. Blots were detected by ECL plus Western Blotting substrate (Pierce) or ECL Bright Quantum (Diagomix) on BlueDevil® autoradiography film (Genesee Scientific) or under the ChemiDoc imaging system (BioRad). Primary antibodies were used at the following dilutions: Pol κ (from Dr. T Nohmi 1/1000), Pol κ (from Sigma HPA012035 1/1000), Chk1 (Abcam ab32531 - 1/1000), Chk1-pS345 (Cell signaling 2348 −1/1000), ORC2 (MBL M055-3 - 1/1000), ORC4 (Transduction laboratories H83120, 1/500) MCM7 (Santa Cruz sc22782 - 1/1000), MCM2 (Abcam ab4461 - 1/2000), Actin (Millipore MAB1501 1/10000), Actinin (Millipore 05-384 - 1/2000), Fibrillarin (Sigma-Aldrich SAB4300633 - 1/1000), p53 (BDSciences 554294 - 1/2000). Where indicated, proteins were quantified using ImageJ software.

### RNA extraction and quantitative PCR Analysis

Total RNA was extracted from cell using RNeasy® kit (Qiagen) according to the supplier’s instructions, then 1μg of total RNA was reverse transcribed into cDNA using Superscript II (Invitrogen) according to the manufacturer’s recommendations. Duplicate quantitative PCR assays were run on StepOne Real Time system from Applied Biosystems with TaqMan Universal Master Mix II and specific probes from assay on demand (Applied Biosystems). Three house-keeping genes *(ACTIN B, YWHAZ, GAPDH)* were also amplified and used as references. The relative amount of each mRNA level was normalized to the control condition and calculated using the ΔΔCT method.

### EDU incorporation, QIBC and Chk1 immunofluorescence

For immunostaining, transfected cell lines used for immunofluorescence were plated on an 8-well chamber slide (Lab-Tek®). When EdU incorporation was performed, it was added 15 or 30 minutes to parallel cultures growing exponentially in culture media containing at final concentration of 10 μM. Fixation was done with a 4 % PFA/PBS solution for 10 min, then permeabilization and saturation step was performed in PBS 1X containing BSA 3% and Triton 0.1X for 30 min (PBS-Triton/BSA). Detection of EdU was performed prior to incubation with the primary antibodies using the Click-iT™ Plus EdU Alexa Fluor™ 488 Imaging Kit according to the manufacturer’s instructions (Thermo-Fisher Scientific). Chk1 primary antibody (Abcam ab32531 - 1/1000) was diluted in PBS-Triton/BSA and incubated 1 hour at room temperature. The 8-well chambers slides were washed three times with PBS 1X and incubated in PBS-Triton/BSA containing fluorescent secondary antibodies at 1/800 (Alexa fluorophores, Life Technologies) for 30 min. Three washes were performed and nuclei were stained with DAPI. For mounting the slides, the Prolong Diamond Antifade reagent (Thermo-Fischer Scientific) was used. 10 to 20 images were acquired randomly. After acquisition, the images were processed for automated analysis with the Cell Profiler image analysis software. DAPI signal was used for segmentation of the nuclei according to intensity threshold, generating a mask which identified each individual nucleus as an individual object. This mask was applied to quantify pixel intensities in the different channels for each individual cell/object. The values quantified for EdU and DAPI staining per cell were graph plotted by dual-parameter (EdU *vs* DNA) generating diagrams in a flow-cytometry-like fashion (QIBC Quantitative-based image Cytometry) for each cell condition. This approach allows the assignment of cells to G1, S or G2/M phases.

### Microscope image acquisition and analysis

Image acquisition of multiple random fields was carried out on a wide field Nikon Eclipse Ni-E microscope equipped with x63 oil immerged objective Nikon Plan Apo 1.4 λ using a C-mos DsQi2 camera driven by NIS-Elements AR software. Fluorescence quantifications were performed with Cell Profiler2.1.1. Images were assembled with Adobe photoshop® and Adobe Illustrator®.

### DNA fiber assay

Cells were pulse-labelled with 50 μM ldU for 20 minutes, washed, then treated with HU (1mM) for 1 hour. After removal of HU and washing, ongoing DNA fibers were labelled in medium containing CldU at 100μM final for 30 minutes. The cells were harvested, lysed in 200 mM Tris-HCl pH 7.4, 0.5% SDS, 50 mM EDTA, and the DNA fibers were spread on glass slides. The slides were incubated with 0.5mg/ml pepsin in 30 mM HCl at 37°C for 20 minutes, the DNA was denatured in 2.5 M HCl for 1 hour and blocked with 1% BSA containing 0.1% Tween 20 in PBS1X. The nucleotide analogues were detected with primary antibodies against CldU (Novus), and IdU (BD Biosciences) and the secondary antibodies anti-rat Alexa Fluor 555 and anti-mouse Alexa Fluor 488 (Thermo Fischer Scientific). Coverslips were mounted on slides using the Prolong Diamond Antifade reagent (Thermo-Fisher Scientific). Images were captured with a Nikon Ni-E microscope and a DS-Qi2 camera equipped with a 20X objective and lengths of CldU tracks were measured with NIS-Elements AR imaging software.

### Statistical analysis

GraphPad Prism version 5.03 for Windows was used for statistical analysis. Differences were considered statistically significant at *P* < 0.05. Data are reported as the medians with 25%-75% interquartile ranges or as the means ± SEM. Results were compared by 2-tailed Student’s *t* test for two groups or a Mann-Whitney nonparametric test as written in legends.

## Acknowledgements

We thank Drs V Bergoglio and S Manenti from the Cancer Research Center of Toulouse for the critical reading of the manuscript and helpful discussions. We thank Dr Takehiko Nohmi (National Institute of Health Sciences, Japan) and Dr Masami Yamada (National Institute of Health Sciences, Japan) for the Rabbit anti-Polκ antibodies. M Dall’Osto is financially supported by the Region Languedoc Roussillon Midi Pyrénées /INSERM fellowship (U1037-R16068BB-Region). Work was supported by funding from INCa-PLBIO 2016, ANR PRC 2016, Labex Toucan and La Ligue contre le Cancer (Equipe labellisée) to JSH, as well as Ligue Régionale grants RAB17004BBA to MJP.

## Author contribution

NV performed the rescue experiments with the GFP-Pol Kappa vector. MJP, MD and LP performed experiments. MD and MJP conceived the study design. MD and LP defined experimental approaches. MD carried out data analysis with the supervision of MJP. MD, MJP and JSH wrote the manuscript. JSH and MJP acquired the fundings.

## Conflicts of Interest

The authors have no financial or other conflict of interest to declare.

## Supplementary materiel and methods

Cells were lysed in a buffer containing 50 mM Tris pH 7.5, 300 mM NaCl, 1% Triton, 5 mM EDTA, 1 mM DTT, supplemented with inhibitors. Cell lysates were cleared by centrifugation for 10 min at 10 000 rpm. Cell extracts were boiled in loading buffer (Biorad 4X laemmli sample buffer) added with DTT 0.1M.

**Supplemental figure 1:**
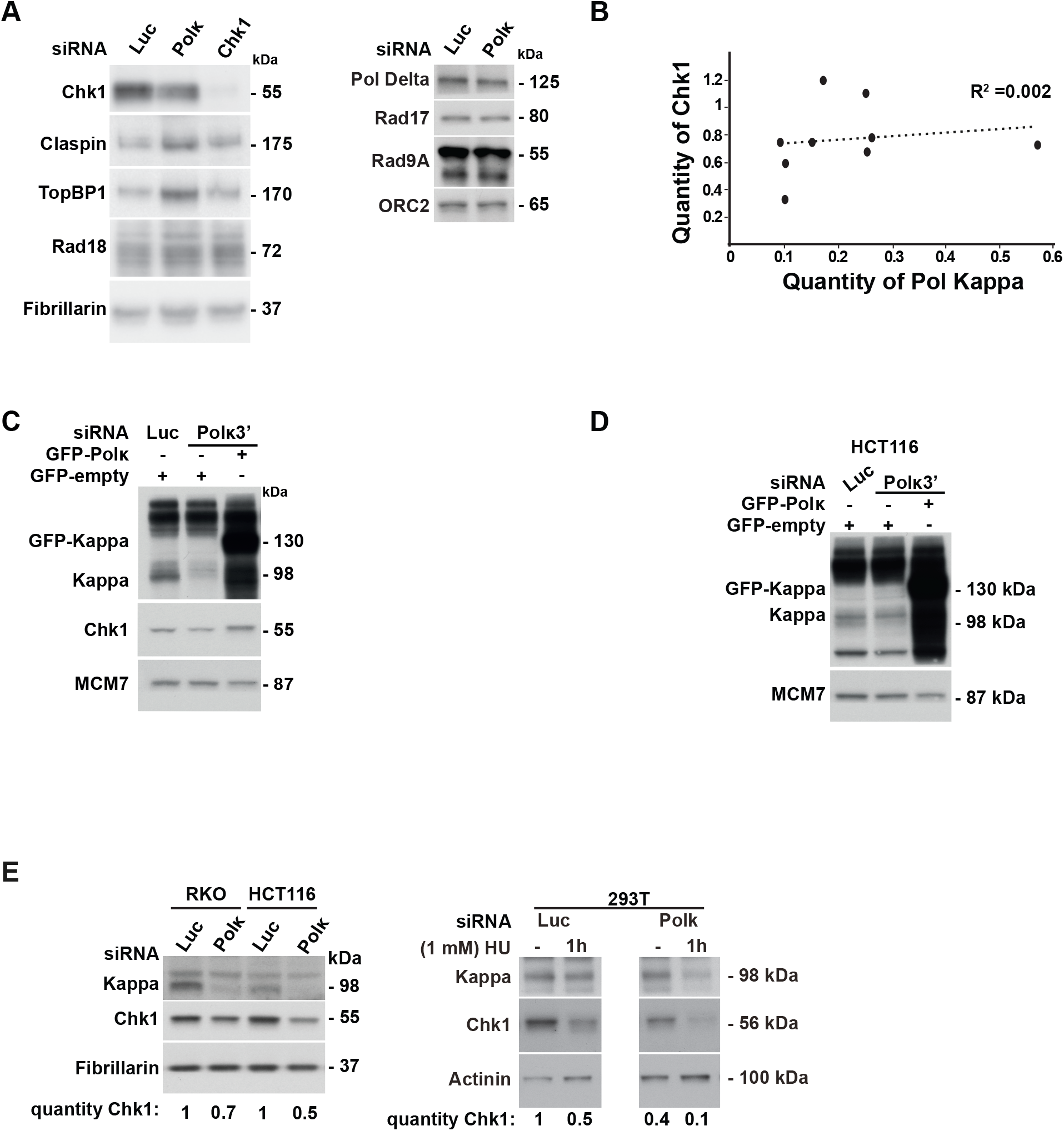
Chk1 protein level is reduced in the nucleus of mammalian cells depleted for Pol κ. (A) Western Blot analysis of nuclear extracts from MRC5-SV cells transfected with a control siRNA (Luc), siRNA targeting the coding sequence of Pol κ (Polκ) or Chk1 (Chk1). Immunodetected proteins are indicated in the figure. Fibrillarin and ORC2 are shown as loading controls. (B) Relative Chk1 or Pol Kappa protein levels in siPol κ whole cell extracts of MRC5-SV cells were normalized to siLuc condition (quantity of Chk1, quantity of Pol Kappa respectively). Data from 9 independent experiments are plotted. The regression curve (dashed line) and R-square are shown. Western Blot analysis of Pol κ and GFP-Pol κ in nuclear extracts of MRC5-SV (C) and HCT116 cells (D), 48h after co-transfection with a control siRNA (Luc) or targeting the 3’UTR of Pol κ (Polκ3’) with a vector expressing either GFP-empty or GFP-Pol κ (GFP-Polκ). MCM7 is shown as a loading control. (E) Western blot analysis of Chk1 and Pol κ in RKO, HCT116 and 293T nuclear extracts 48h after transfection with a control siRNA (Luc) or Pol κ si RNA (Polκ). Cells were untreated or treated with 1mM HU for 1h. Fibrillarin or actinin are shown as protein-loading controls. Quantification of Chk1 is relative to siLuc condition.

**Supplemental figure 2:**
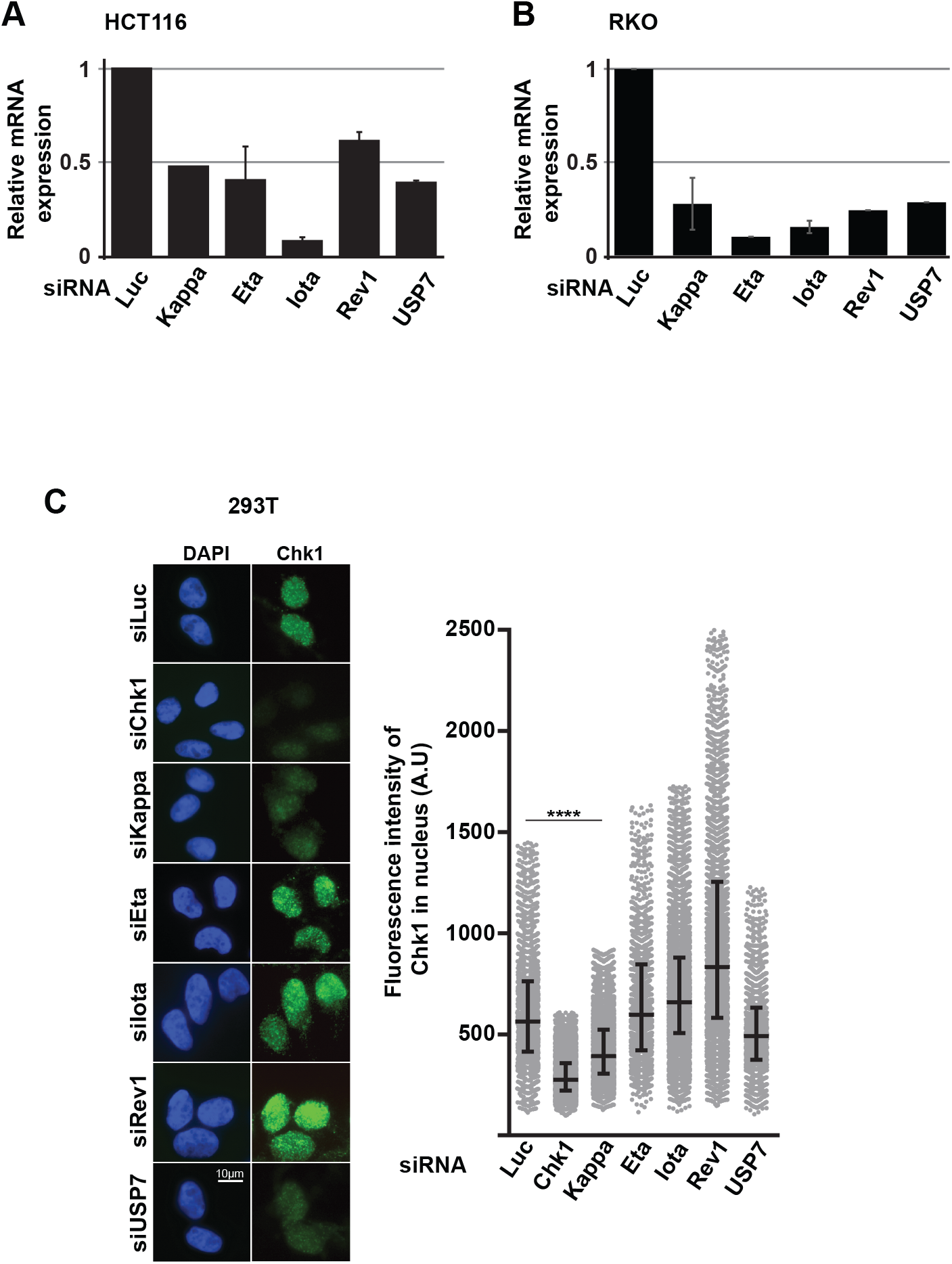
Among the Y-DNA polymerase family, only the Pol κ depletion causes a Chk1 nuclear drop. Efficiency of mRNA silencing was measured by RT-qPCR in HCT116 (A) and RKO (B) cells. The mRNA quantified is the one targeted by the siRNA indicated on the figure. Relative expressions were normalized to siLuc condition. (C) Representative images and quantification of Chk1 immunostaining (green) in 293T cells transfected with the indicated siRNAs and DNA was stained with DAPI. The fluorescence intensity of Chk1 was quantified in each nucleus. Medians with 25% and 75% interquartile ranges were represented (****p<0.0001; Mann-Whitney test). Scale bar, 10 μm. A.U = arbitrary units.

**Supplemental figure 3:**
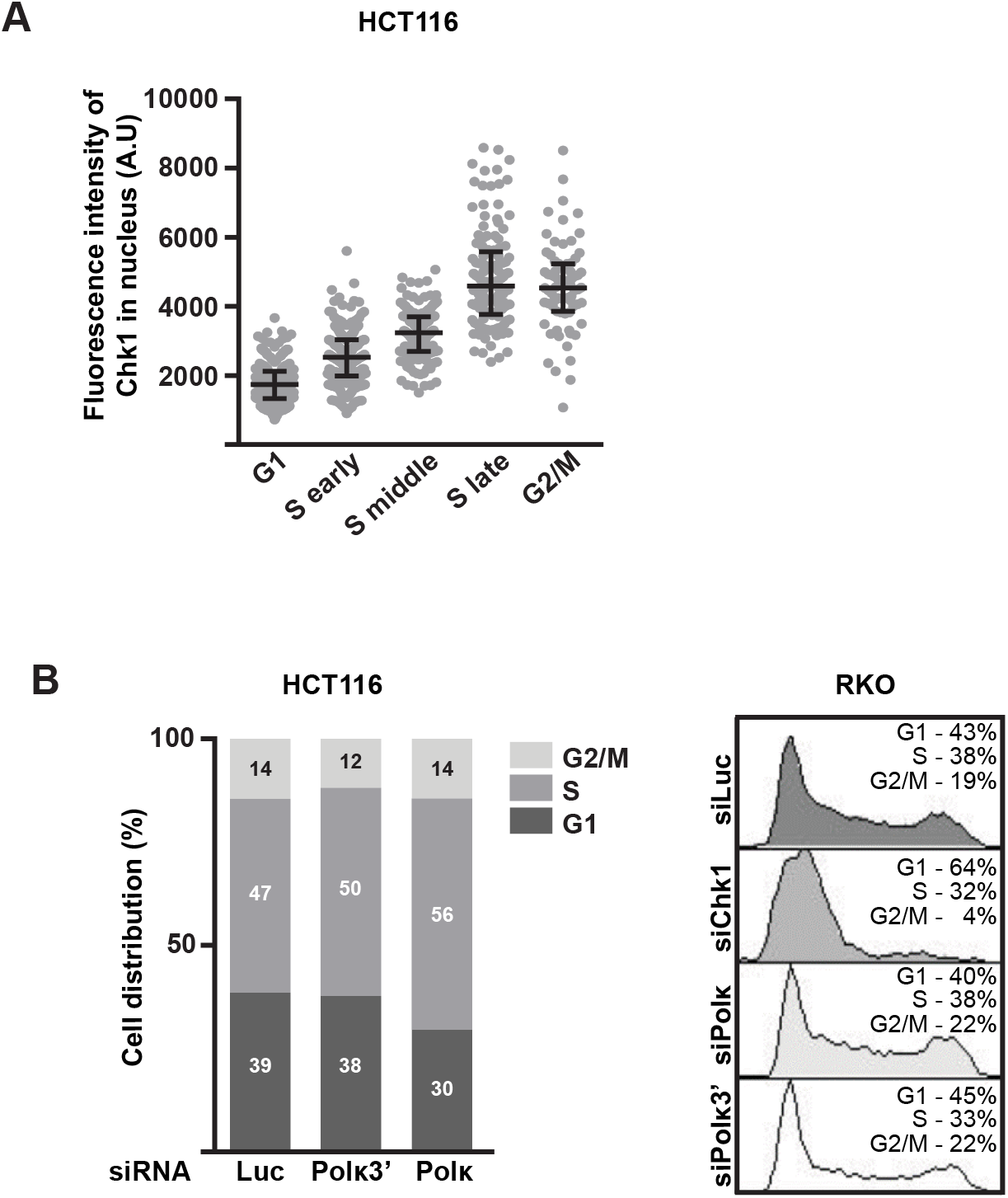
The Pol κ-dependent Chk1 downregulation is not due to cycle arrest in G1. HCT116 or RKO cells were transfected with control siRNA (Luc), siRNA targeting the 3’UTR of Pol κ (Polκ3’), the coding sequence of Pol κ (Polκ) or Chk1 (Chk1). 48h after transfection, asynchronous cells were pulse-labelled with EdU for 30 min (10μM) and processed as in figure 3. (A) Quantification of Chk1 immunostaining all along the cell cycle in HCT116 cells transfected control siRNA. A.U = arbitrary units. (B) No G1 accumulation in Pol κ depleted cells. Cell cycle distribution from QIBC analysis in HCT116 cells *(left)* or from FACS analysis in RKO cells *(right)* after transfection with indicated siRNAs. Percentage of cells in each phase is indicated in the bar graphs or the FACS profiles.

**Supplemental figure 4:**
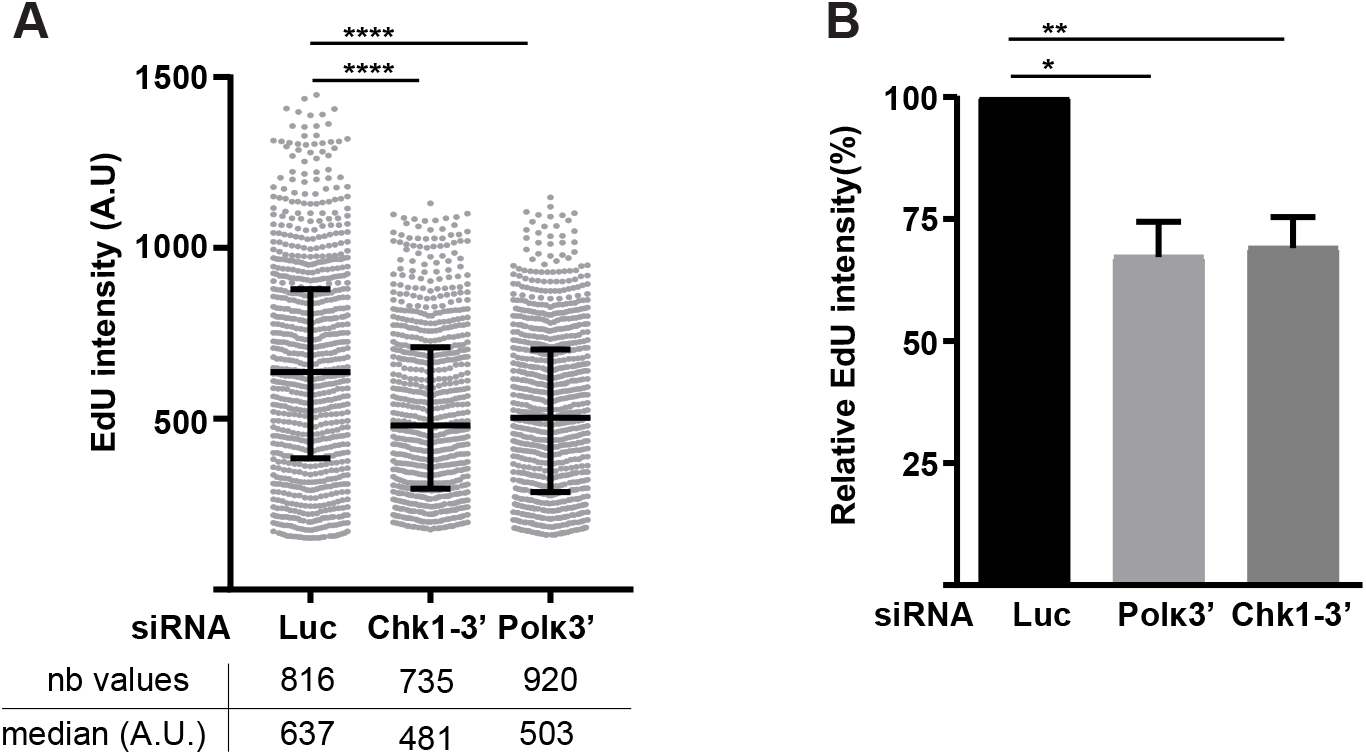
Pol κ is required to maintain the global replication rate. (A) Cells were transfected with control siRNA (Luc), siRNA targeting the 3’UTR of Chk1 (Chk1-3’) or Pol κ (Polκ3’). 48h after transfection, asynchronous cells were pulse-labelled with EdU for 30 min (10μM). Nuclear EdU intensities were quantified in S phase nuclei. Medians with 25% and 75% interquartile ranges are represented. (****p<0.0001; Mann-Whitney test) A.U = arbitrary units. (B) EdU intensity relative to control conditions is determined in 3 independent experiments and the mean (± SEM) of medians relative to siLuc condition is presented. (*p<0.05; **p<0.01; t-test).

## Notes

### Competing Interest Statement

The authors have declared no competing interest.

